# Application of genome-scale models of metabolism and expression to the simulation and design of recombinant organisms

**DOI:** 10.1101/2023.09.13.557522

**Authors:** Omid Oftadeh, Vassily Hatzimanikatis

## Abstract

The production of recombinant proteins in a host using synthetic constructs such as plasmids comes at the cost of detrimental effects such as reduced growth, energetic inefficiencies, and other stress responses, collectively known as metabolic stress. Increasing the number of copies of the foreign gene increases the metabolic load but increases the expression of the foreign protein. Thus, there is a trade-off between biomass and product yield in response to changes in heterologous gene copy number. This work proposes a computational method, rETFL (recombinant Expression and Thermodynamic Flux), for analyzing and predicting the responses of recombinant organisms to the introduction of synthetic constructs. rETFL is an extension to the ETFL formulations designed to reconstruct models of metabolism and expression (ME-models). We have illustrated the capabilities of the method in four studies to (i) capture the growth reduction in plasmid-containing *E. coli* and recombinant protein production; (ii) explore the trade-off between biomass and product yield as plasmid copy number is varied; (iii) predict the emergence of overflow metabolism in recombinant *E. coli* in agreement with experimental data; and (iv) investigate the individual pathways and enzymes affected by the presence of the plasmid. We anticipate that rETFL will serve as a comprehensive platform for integrating available omics data for recombinant organisms and making context-specific predictions that can help optimize recombinant expression systems for biopharmaceutical production and gene therapy.

## Introduction

Recombinant protein expression involves the transfer of heterologous genes into a prokaryotic or eukaryotic host organism. The foreign genes are delivered to the host using an engineered DNA molecule called a vector. There are several types of vectors, but the plasmid is the most common. A plasmid can carry functional genes and provide its host with selective advantages, such as antibiotic resistance. The presence of the plasmid in the host can also trigger metabolic stress responses such as a reduction in growth (1, 2), an increase in maintenance energy (3, 4), and the emergence of overflow metabolism (5, 6). Such stress responses are referred to as plasmid metabolic load. The plasmid load depends on several factors, including copy number, number of genes on the plasmid, and strength of the promoters on the plasmid.

Most of these approaches have focused on simulating the plasmid load in *E. coli* as the most widely used host for the expression of recombinant proteins. Peretti and Bailey reconstructed a whole-cell kinetic model that included key cellular processes such as DNA replication, mRNA transcription, and protein translation (7). However, as the kinetic parameters and mechanisms for many biological reactions are unknown, they greatly simplified the cellular processes. da Silva and Bailey developed a theoretical model to calculate the plasmid effect on biomass yield when the additional energy and material requirements caused by the plasmid are known (8). Bentley et al. developed a structured kinetic model to investigate the relationship between growth rate and the level of heterologous protein expression (9). To this end, they included separate reactions for plasmid-related DNA, mRNA, and protein synthesis in the model. Özkan et al. used constraint-based optimization to capture the plasmid load (10). They used a stoichiometric model to represent cell metabolism under the steady-state assumption, where a single reaction was added to represent the plasmid-related energy and material requirements. Experimental fluxomic data were used to constrain the fluxes in the central metabolism, and an optimization problem was solved to find the other fluxes. In another study, Ow et al. integrated a lumped reaction that accounts for plasmid requirements into a genome-scale metabolic model (GEM) (4). They explored different objective functions to find the cellular objective that was most consistent with the experimental data. Recently, Zeng and Yang integrated empirical constraints into the E. coli GEM to account for foreign protein expression and plasmid maintenance requirements (11).

Metabolism and Expression models (ME-models) are constraint-based models that simulate cellular metabolism and expression (12-14). Reconstruction of an ME model starts with a GEM representing metabolism, and additional constraints are incorporated to account for expression. Expression and Thermodynamics-enabled Flux (ETFL) is a mixed-integer linear formulation for the reconstruction of ME models (14-16). The previous formulations of ME-models were nonlinear and required special quad-precision solvers (12, 13). In contrast, ETFL is a linear formulation that can be solved with standard double-precision solvers. dETFL is an extended version of ETFL that considers temporal dynamics of extracellular metabolite concentrations and enzyme abundances (15). Recently, we have extended the ETFL formulation to the study of eukaryotic organisms. To this end, we enabled the implementation of multiple RNA polymerases and ribosomes and accounted for the compartmentalized expression systems in eukaryotes. We also improved the parameterization of the ETFL models by correcting for growth-associated maintenance (GAM) and allocating a limited proteome fraction to metabolic and expression-related enzymes. We used the extended ETFL formulation to reconstruct the first ME model for *Saccharomyces cerevisiae*, yETFL (16).

This work presents an updated ETFL model for *E. coli*, ecETFL, by improving the model parameters, including GAM and resource allocation. We also extend the ETFL formulation to allow the simulation of recombinant cells. The proposed formulation, called rETFL, allows the user to include new genes in the model and to integrate new constraints for the allocation of expression resources to plasmid-related macromolecules. We used rETFL to simulate the plasmid load for different plasmids in *E. coli*. The explicit representation of individual enzymes in rETFL allows the investigation of enzymes that are more affected by the presence of the plasmid. Furthermore, rETFL allows the mechanistic investigation of different transcriptomic and proteomic perturbations in recombinant cells.

## Results and Discussion

### Updated *E. coli* ETFL model

In addition to the 1366 metabolic genes from the FBA model, an updated *E. coli* ETFL model, ecETFL, has 69 genes encoding RNA polymerase, ribosomal RNAs, and ribosomal peptides. Since the transcription elongation rate is faster for stable RNA (sRNA) in *E. coli*, we implemented two RNA polymerases (Methods): (i) the faster RNA polymerase with an elongation rate of 85 nucleotides/second, which is associated with rRNAs and tRNAs; and (ii) the slower RNA polymerase with an elongation rate of 45 nucleotides/second, which is associated with the other genes. One ribosome is implemented to translate all mRNAs into proteins. The model includes 1128 metabolic enzymes catalyzing 2007 reactions (Table 1).

**Table 1:** Properties of ecETFL.

|  |  |
| --- | --- |
| Growth upper bound ($\bar{\mu}$) | 1.5 h<sup>-1</sup> |
| Number of bins (N) | 128 |
| Resolution ($\bar{\mu}/N$) | 0.0117 h<sup>-1</sup> |
| Number of species |  |
| - Metabolites | 1809 |
| - mRNAs | 1432 |
| - Peptides | 1432 |
| - rRNAs | 3 |
| Number of enzymes |  |
| - Metabolic enzymes | 1128 |
| - RNA polymerases | 2 |
| - Ribosomes | 1 |
| Number of reactions |  |
| - Metabolic | 1543 |
| - Transport | 733 |
| - Exchange flux | 330 |
| - Transcription | 1435 |
| - Translation | 1432 |
| - Complexation | 1131 |
| - Degradation | 2566 |
| Thermodynamic data |  |
| - Number of metabolites $\Delta G'^{\circ}_f$ | 1737 |
| - Number of reactions $\Delta G'^{\circ}_r$ | 1787 |

As a benchmark for ecETFL, we simulated the growth rate at different glucose uptake rates (Figure 1A). Initially, growth increased linearly with increasing the uptake rate. In this part, growth is limited by substrate availability, and both the FBA and ecETFL models were able to capture the experimental data. However, as the cellular expression capacity is limited, the growth reached a plateau that could not be further increased by increasing the uptake. While FBA failed to capture the shift from substrate-limited to protein-limited growth, ecETFL predicted that growth would reach a maximum in accordance with the experimental data (Figure 1a). The observed maximum growth rate of *E. coli* in the minimal medium was 0.61 h^-^ (17), whereas ecETFL predicted a maximum growth rate of 0.67 h^-1^. The agreement between the predicted and measured maximum growth rate shows that the updated ecETFL model improves upon the previous ME-models for *E. coli* (12, 14), as these models captured the maximum growth rate with a significant deviation from the experimental observations.

**Figure 1:**
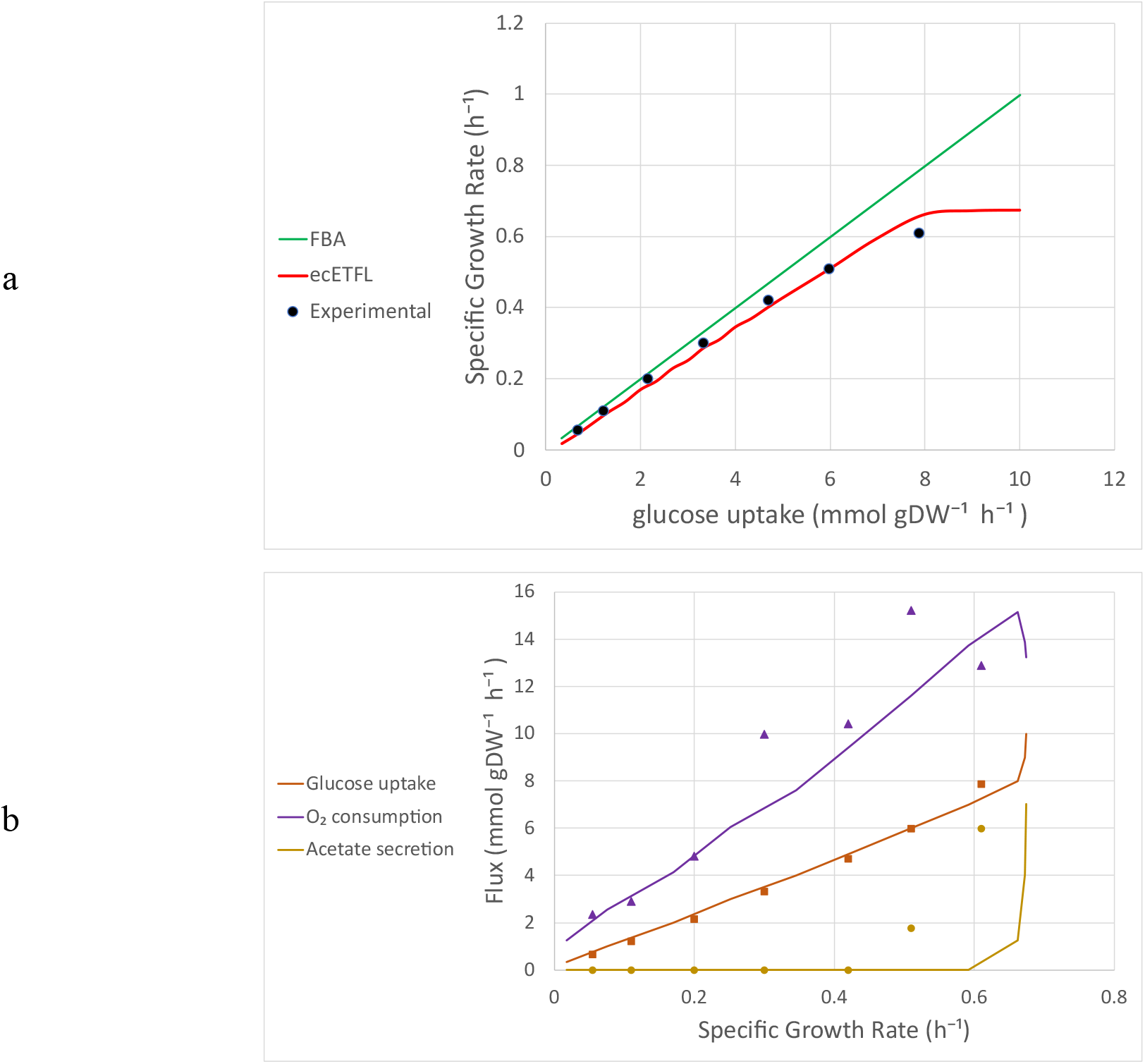
Benchmarking ecETFL against experimental data. **a** the simulation of maximum growth rate (h^-1^) at different glucose uptake rates (mmol gDW^-1^ h^-1^). ecETFL captured that the growth rate plateaued at high glucose uptakes due to the limited enzymatic capacities. The model predicted a maximum growth rate of 0.67 h^-1^, close to the experimental maximum growth rate of 0.61 h^-1^. **b** the simulation of overflow metabolism in *E. coli*. ecETFL predicted a shift in metabolic fluxes of acetate secretion, glucose uptake, and oxygen consumption after a critical growth rate of 0.58 h^-1^. The model predictions were in qualitative agreement with the experimental data, which showed the oxygen consumption decrease and the emergence of acetate production after the growth rate of 0.42 h^-1^. The experimental data were taken from Vemuri et al. (17).

Overflow metabolism is a shift from pure respiration to a combination of respiration and fermentation observed in fast-growing cells (18-20). This shift results in seemingly suboptimal secretion of fermentation byproducts, which could otherwise be incorporated into the biomass. One hypothesis is that overflow metabolism occurs due to the limited capacity of the enzymes involved in respiration and redox balance (21-23). As the ETFL formulation considers the limited enzymatic capacity through the catalytic constraints, we investigated the ability of ecETFL to capture overflow metabolism in E. coli as a further test of the quality of the model (Figure 1b). At growth rates above a critical growth rate, which is strain specific but estimated to be around 0.42 h^-1^, *E. coli* cells secrete acetate while consuming oxygen, known as overflow metabolism in *E. coli*. ecETFL predicted the shift in metabolic fluxes at high growth rates, albeit delayed with respect to the experimental data. The model captured the decrease in acetate secretion and oxygen consumption at growth rates above 0.58 h^-1^. The same delay in the predicted onset of overflow metabolism was observed in *Saccharomyces cerevisiae* using yETFL (16). In that paper, we discussed that improvements such as the inclusion of regulatory constraints or the integration of more growth-dependent parameters could further reconcile model predictions and experimental data (16).

### Quantifying the allocation of resources to the expression of heterologous genes

rETFL has three additional parameters that quantify the allocation of resources to heterologous gene expression. The first two parameters, 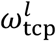and 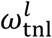, are phenomenological parameters that determine the basal level of RNA polymerases and ribosomes, respectively, allocated for the heterologous gene *l* expression. 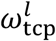characterizes the availability of the promoter of the gene *l* and the affinity of RNA polymerase to this promoter. Similarly, 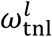represents the affinity of ribosomes to the mRNA *l*. The third parameter, *φ*_*h*_, represents the fraction of the heterologous proteins taking their share from the metabolism- and expression-related (ME-) enzymes (see Methods for more details). Since ME enzymes synthesize biomass building blocks and generate energy for various cellular processes, allocating a higher proportion of the ME enzyme fraction to the heterologous proteins represents a higher metabolic burden (24).

We used data on the fraction of RNA polymerase and ribosome assigned to the plasmid (7) to estimate 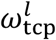and 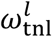at different copy numbers for plasmid pMB1. Table S1 summarizes the estimated values of 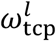and 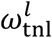. It should be noted that the values of 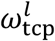 and 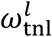might vary subject to different promoters and ribosomal binding sites. We observed that the specific activity of RNA polymerase and ribosome decreased with increasing copy number. We fitted the model to experimental data (7) to estimate *φ*_*h*_. For plasmid pMB1, we obtained a proper fit to the data with *φ*_*h*_= 0.2, implying that 20% of the heterologous proteins recruit the resources allocated to the ME-enzymes.

In addition to the additional requirements for the expression of heterologous genes, plasmid burden manifests itself in increased energy requirements for maintenance (3, 4). As a result, plasmid-containing cells are less energetically efficient than wild-type cells. This increase in global maintenance energy is attributed to plasmid maintenance. ATP maintenance (ATPM) is an ATP hydrolysis reaction added to the model to account for global energy maintenance. The level of ATPM is determined by fitting model predictions to experimental growth (25). For *E. coli*, different levels of ATPM have been reported for different strains of *E. coli* and different versions of GEMs (25, 26). For example, the ATPM is set to 3.15 mmol gDW^-1^ h^-1^ in iJO1366 (26) and 8.39 in iAF1260 (25) for wild-type *E. coli*. To account for the reduced energetic efficiency caused by the introduction of the plasmid, we estimated the ATPM to be 15 mmol gDW^-1^ h^-1^ by fitting the model to the experimental growth in recombinant *E. coli* containing pMB1 (7).

#### The plasmid impact on growth rate

We used ecETFL and the fitted parameters to simulate the maximal growth of recombinant *E. coli* containing different copy numbers of pMB1 (Figure 2a). At low copy numbers, where a smaller fraction of resources was allocated to heterologous synthesis, the metabolic load was dominated by energy requirements for plasmid maintenance. As copy numbers increased, the fraction of resources allocated to plasmids also increased, and the metabolic burden was mainly due to the additional requirements for the synthesis of plasmid-related macromolecules. The recombinant ecETFL also predicted the relative heterologous protein production according to the experimental data (Figure 2b). Heterologous protein production increased non-linearly with increasing copy number and reached a maximum where no more resources could be allocated to the plasmids.

**Figure 2:**
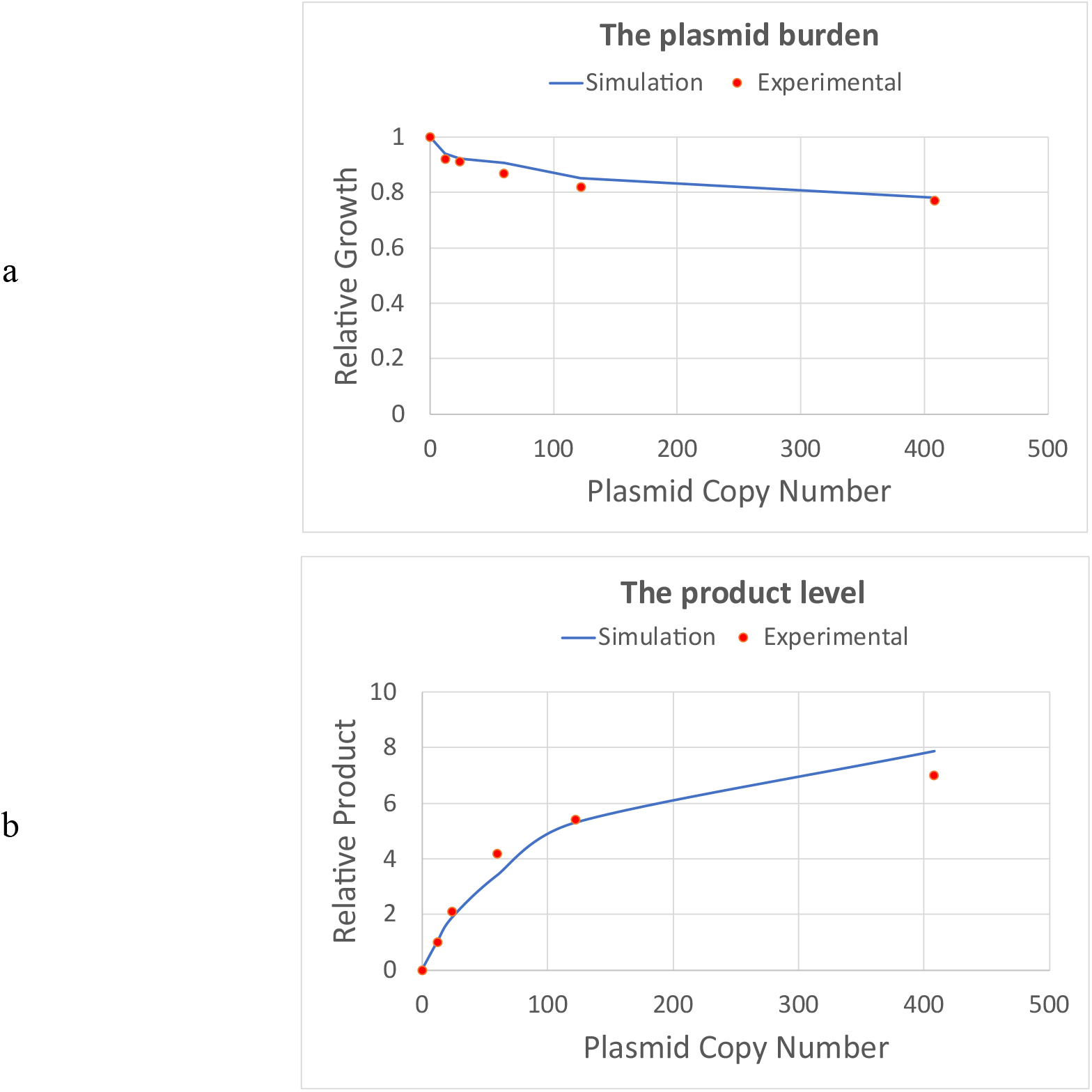
Relative growth and product formation as a function of pMB1 copy number. **a** The presence of the plasmid exerts a metabolic burden on the host due to extra resource requirements and energetic inefficiency. The metabolic burden manifests as decreased growth rate. Increasing the plasmid copy number adversely affects biomass yield. **b** The amount of heterologous protein produced from the plasmid, i.e., the product, increases with increasing the copy number. However, the increase in the product level is nonlinear and reaches a maximum due to the saturation of expression enzymes.

### The impact of plasmid copy number on biomass and product yields

The heterologous protein may benefit the host by providing a novel metabolic function or enhancing an existing capacity. Applying evolutionary pressure can translate such benefits into selective advantages. For example, appropriate evolutionary pressure stimulates higher heterologous protein production in the host. For example, if the product protein confers antibiotic resistance, adding antibiotics to the medium can further stimulate product production. We simulated the stimulated product production using a multi-objective problem with two objective functions, i.e., maximizing growth and maximizing heterologous protein production:

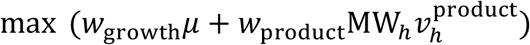

with *w*_growth_ and *w*_product_ denoting arbitrary weights assigned to the objectives such that *w*_growth_ + *w*_product_ = 1, *μ* is the specific growth rate, and MW_*h*_ and 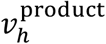 represent the molecular weight and the production rate of the heterologous protein, respectively. We explored the trade-off between the two objectives by assigning different weights (Figure 3). As expected, for *w*_growth_ = 1, the minimum product yield increased with increasing the copy number. If the product was not beneficial to the host, increasing the copy number increased the product yield, but at the expense of decreasing the biomass yield.

**Figure 3:**
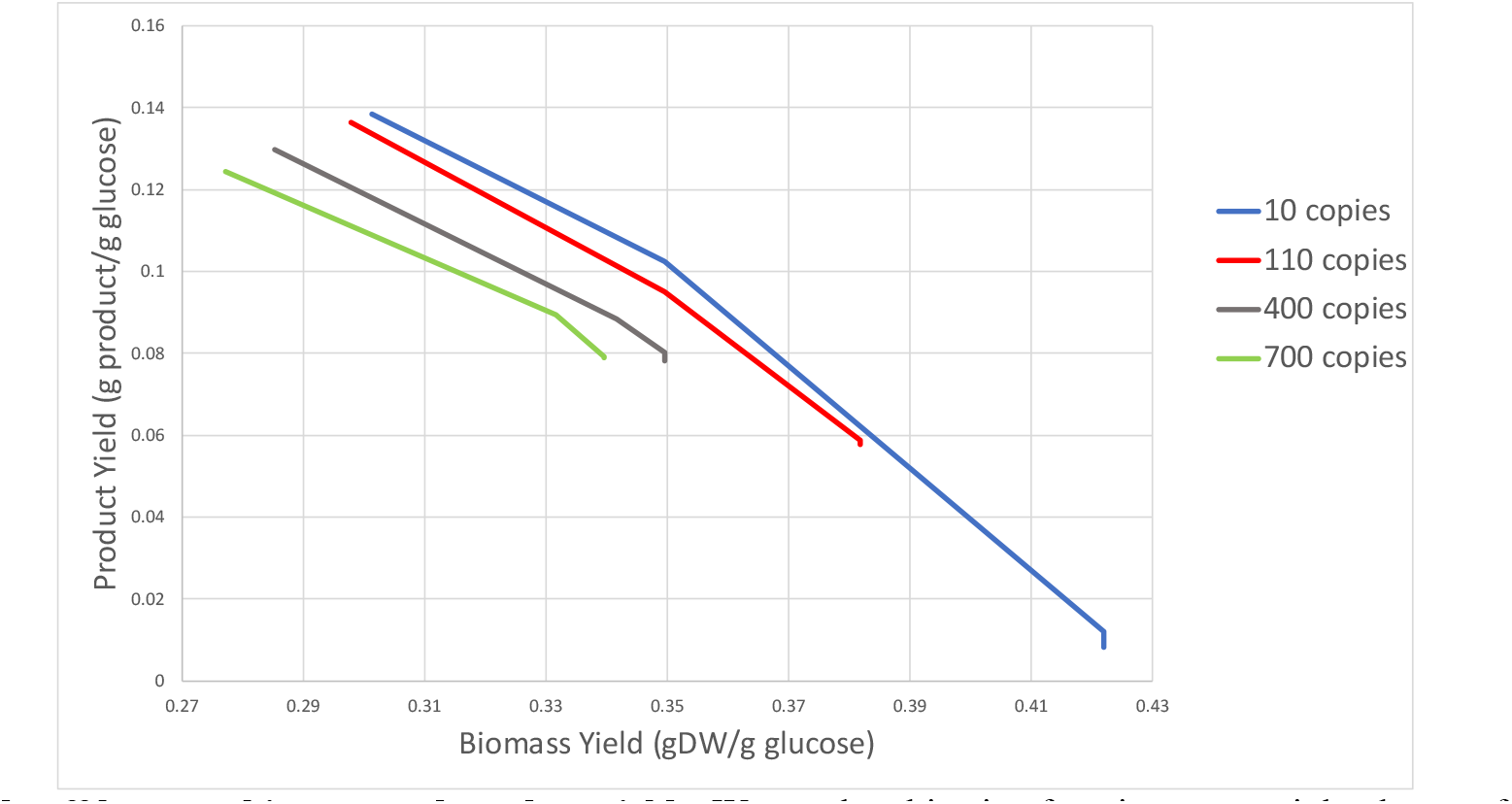
Trade-off between biomass and product yields. We set the objective function as a weighted sum of growth rate and heterologous protein concentration. We changed the objectives’ weights subject to different plasmid copy numbers to explore the Pareto front. An increase in the copy number raised the minimum product yield but at the expense of reducing the biomass yield. On the other hand, an increase in the copy number decreased the maximum product yield due to allocating more resources to plasmid-related RNA and DNA. The most optimal solutions were obtained when the copy number was low, but the product production was motivated by the objective function.

On the other hand, if product production was the sole cellular objective, i.e., *w*_product_= 1, increasing the copy number reduced the maximum product yield due to the additional requirements for plasmid-related RNA and DNA synthesis. Indeed, when the objective function stimulated the product production at low copy numbers, higher product yields were achieved than when the production was enforced by increasing the copy number. Our results suggest that the stimulated product production, e.g., by exerting proper selective pressure, is more efficient than increasing the copy number because higher product and biomass yields are achieved.

#### The impact of plasmid on consumption and secretion fluxes

For this study, we used rETFL to simulate the metabolic burden of plasmid pOri2 and its effect on acetate secretion and oxygen consumption. Like pMB1, pOri2 genes are transcribed under the lac promoter. Therefore, we used the same values for RNA polymerase and ribosome affinities for the plasmid genes, i.e., 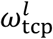and 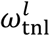, as was used for pMB1 (Table S1). We varied the fraction of resources allocated to the plasmid-related proteins, *φ*_*h*_ and the ATPM so that the model fits the experimental growth of *E. coli* containing pOri2 (6). The estimated values of *φ*_*h*_ and the ATPM were, respectively, 30% and 30 mmol gDW^-1^ h^-1^. Interestingly, the estimated value of ATPM obtained was close to that obtained in Zeng and Yang using a phenomenological model (11). The ATPM found for pOri2 was significantly higher than pMB1 (15 mmol gDW^-1^ h^-1^), indicating that pOri2 is energetically less efficient.

We then used ecETFL to compare the model predictions for oxygen consumption and acetate secretion with the experimental data in the wild-type and plasmid-containing organisms (Table 2). The model captured the impact of the plasmid on the exchange fluxes in agreement with the experimental observations. Notably, the model predicted acetate production in the plasmid-containing *E. coli*, whereas no acetate was produced in the wild-type organism.

**Table 2:** the predicted and experimental growth rate, oxygen consumption, and acetate secretion in wild-type *E. coli* (copy number = 0) and recombinant *E. coli* containing pOri2 (copy number = 410). Glucose uptake was constrained by an upper bound of 5.2 and 6.3 mmol gDW^-1^ h^-1^, the values measured in the wild-type and recombinant cell, respectively. The experimental data were obtained from Wang et al. (6). Abbreviations: Ex.: Experimental measurement; Mod.: Model prediction.

| Copy number | Glucose uptake (mmol gDW<sup>-1</sup> h<sup>-1</sup>) | Growth (h<sup>-1</sup>) |  | Acetate secretion (mmol gDW<sup>-1</sup> h<sup>-1</sup>) |  | Oxygen uptake (mmol gDW<sup>-1</sup> h<sup>-1</sup>) |  |
| --- | --- | --- | --- | --- | --- | --- | --- |
|  |  | Mod. | Ex. | Mod. | Ex. | Mod. | Ex. |
| 0 | 5.2 | 0.44 | 0.46 | 0 | 0 | 11 | 11 |
| 410 | 6.3 | 0.29 | 0.29 | 5.5 | 4.4 | 13.2 | 12.2 |

### Proteome comparison in the wild-type and recombinant organisms

By explicitly simulating the expression of individual proteins, we were able to use rETFL to evaluate the differences in the proteomes of wild-type and recombinant *E. coli*. In the recombinant organism, part of the proteome is allocated to the heterologous proteins, limiting the resources available to the native proteins. We compared the levels of several enzymes in wild-type and recombinant *E. coli*. We calculated a normalized expression score (*s*_*j*_) for each protein according to this formula:

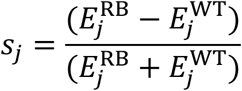

where 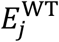and 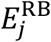are the concentrations of enzyme *j* in the wild-type and recombinant organisms, respectively. If the enzyme *j* is upregulated due to the presence of the plasmid, *s*_*j*_ is positive, and if the enzyme *j* is downregulated, *s*_*j*_ is negative (Figure 4). Out of the 1131 enzymes included in the model, 778 enzyme concentrations remained almost unaffected by the presence of plasmid, i.e., −0.1 < *s*_*j*_ <0.1. Due to the allocation of cellular resources to the heterologous proteins, most of the remaining enzymes were slightly downregulated, including 251 enzymes with −0.3 < *s*_*j*_ < −0.1. We found that 34 enzymes were highly upregulated, i.e., 0.5 < *s*_*j*_, and 29 were highly downregulated, i.e., *s*_*j*_ < −0.5.

**Figure 4:**
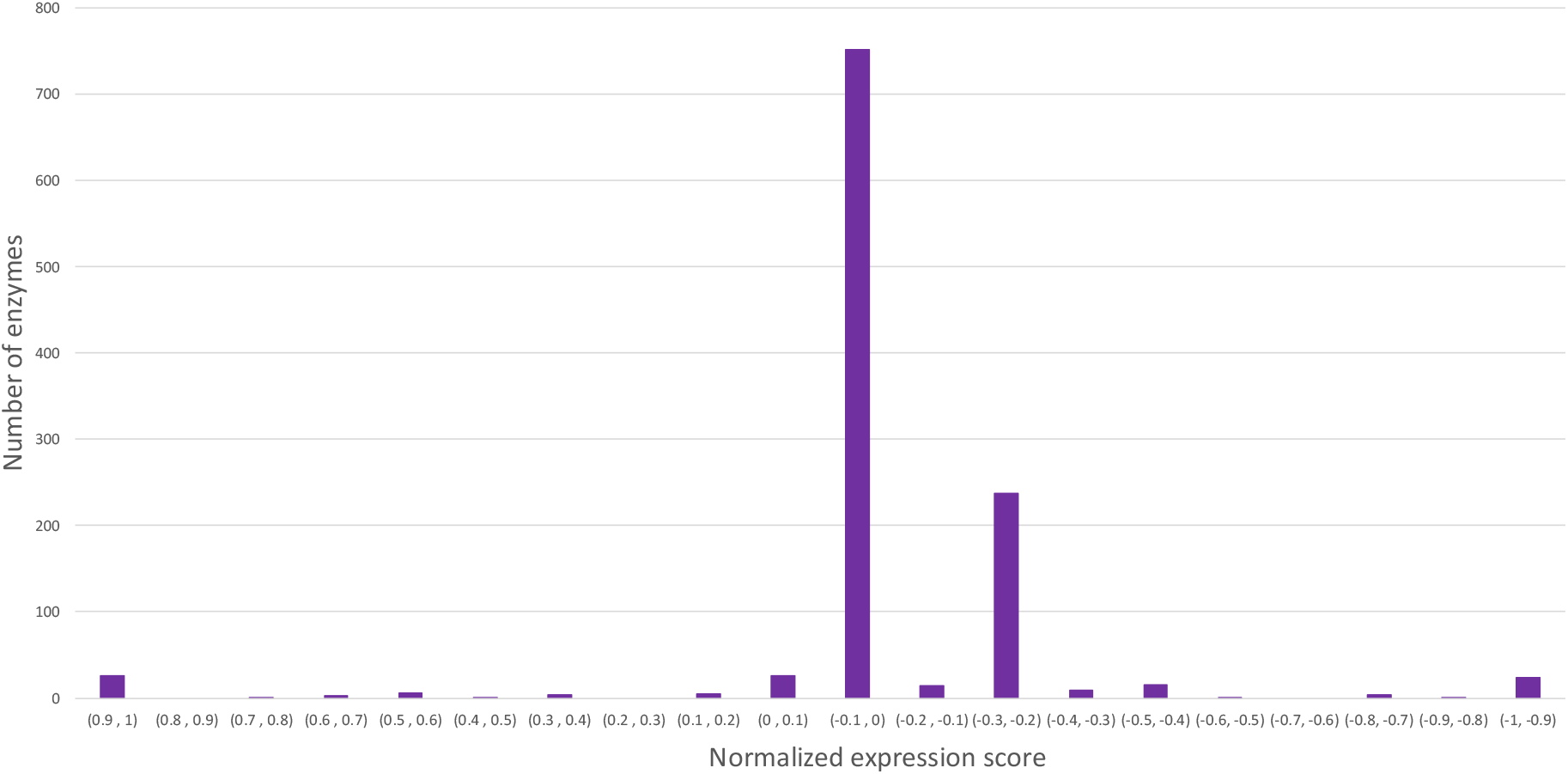
Normalized comparison of expression of different enzymes in wild-type and recombinant *E. coli*. For the normalized expression scores close to 0, the expression of the enzymes was not affected by the plasmid. The positive and negative scores reflect up- and downregulation after inserting the plasmid. While most of the enzymes were unaffected by the plasmid, a total number of 34 and 29 enzymes were highly up- and down-regulated, respectively. We assumed an enzyme expression is highly up- or downregulated, respectively, if the normalized expression score was more than 0.5 or less than - 0.5. Comparing the turnover number (*k*_cat_) and molecular weight of the enzymes with significant changes in their expression, we showed that the enzymes upregulated in recombinant *E. coli* are more mass efficient than the enzymes downregulated.

The maximum catalytic capacity of an enzyme can be represented as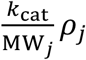, where *ρ*_*j*_ is the mass concentration. As a result, for larger values of 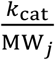, the cell requires smaller amounts of enzymes to achieve the same catalytic capacity. We calculated the average 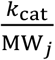to be 3.68 mol g^-1^ min^-1^ for the 34 enzymes upregulated in the recombinant *E. coli*, significantly higher than 0.22 mol g^-1^ min^-1^, the average 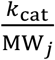for the 29 downregulated enzymes. This implies that the recombinant organism synthesizes enzymes with higher mass efficiencies under more limited resource availability at the expense of switching to a suboptimal metabolism.

## Conclusion

In this work, we presented rETFL, an extension of the ETFL formulation and code to simulate the expression of heterologous genes in recombinant organisms. To this end, we extended the ETFL formulation to account for the allocation of cellular resources and expression machinery to plasmid-related activities. The new formulation allows us to account for the energetic burden imposed by the plasmid by modifying ATP maintenance. We demonstrated that rETFL could capture the plasmid burden and heterologous protein production in recombinant *E. coli*. We also simulated the change in reaction fluxes due to the presence of the plasmid in agreement with the experimental observations without directly constraining the fluxes as in the previous constraint-based formulations of the plasmid burden (4, 10).

rETFL allows the integration of different omics data, including transcriptomics, proteomics, and metabolomics. Since the ETFL models can be readily developed for both prokaryotic and eukaryotic organisms, rETFL can be used to simulate recombinant protein expression in different hosts. Furthermore, like the original ETFL formulation, rETFL can be extended to dynamic settings to capture time-dependent evolutions (15). The mechanistic representation of the expression of individual enzymes in rETFL allows us to reveal the specific pathways and enzymes affected by plasmids. rETFL is available as open-source code for generating and analyzing models of recombinant organisms. We envision that rETFL can be a versatile tool to simulate recombinant organisms and propose metabolic and protein engineering strategies to design optimal hosts for biotechnological applications. In addition, rETFL can simulate and support other types of genetic interventions, such as gene therapies in humans and animals.

## Methods

### Data Collection

The most recent version of iJO1366 was obtained from the BiGG database (27). The essential metabolites to produce 1 gram of biomass were taken from the growth reaction and divided into different types, including amino acids, nucleoside triphosphates, deoxynucleoside triphosphates, lipids, peptidoglycans, lipopolysaccharides, ions, and cofactors. The percentage of different macromolecules in the biomass was then calculated. Sequences of peptides and mRNAs were obtained from the KEGG database (28). The functions from GECKO (29) were used to obtain the turnover numbers (*k*_cat_s). The composition and stoichiometry of the enzymes were obtained from a previous ME-model for *E. coli* (12).

### Updating the *E. coli* ETFL model

The *E. coli* ETFL model presented here, i.e., ecETFL, is improved in three main aspects. First, we incorporated an additional constraint to determine the maximum proteome fraction allocated to the ME-enzymes, as previously done for *Saccharomyces cerevisiae* (16). The latest whole-cell proteomics data for *E. coli* was obtained from PaxDB to calculate the fraction of the ME-enzymes (30). Second, we modified the GAM to avoid double counting the energy requirements for peptide synthesis. According to the biomass reaction in iJO1366, ∼5.2 mmol of amino acids are required to produce 1 gram of biomass. We know 3 mmol of ATP are consumed to attach an amino acid to a peptide chain, including 1 mmol ATP for the tRNA charging and 2 mmol ATP for the amino acid assembly (14, 16). In total, the energetic requirement for peptide synthesis is 3 ×5.2= 15.6 mmol gDW^-1^ of ATP, which was removed from the GAM. Third, we integrated more enzymes into the model such that the number of enzymes in ecETFL is 1131, compared to 562 enzymes in the previous *E. coli* ETFL model. Recently, we extended the ETFL formulation to account for multiple RNA polymerases and ribosomes (16). Like other bacteria, *E. coli* has only one type of RNA polymerase. However, it is observed that its RNA polymerase transcribes the sRNAs much faster than the mRNAs (31). We used the extended ETFL formulation to define two types of RNA polymerases in ecETFL with identical compositions but different catalytic efficiencies. The faster RNA polymerase was associated with the sRNAs, and the slower one with the mRNAs.

### Extending the formulation of ETFL

#### Expression

The original ETFL formulation simulates cell behavior under the optimality assumption where growth is maximized. This means that the ETFL models, like other similar models, could only predict the synthesis of proteins that contribute to the growth of the organism. Such models do not predict the synthesis of proteins that are not beneficial for growth because in this way, a higher fraction of the cellular protein content could be allocated to proteins with a positive contribution to growth. However, the cell could produce gratuitous proteins that have no function in the cell (32). Similarly, heterologous proteins transferred into a host often do not have a positive impact on cellular activity (33). To allow the ETFL formulation to account for the expression of nonfunctional proteins, we incorporated the following two constraints:

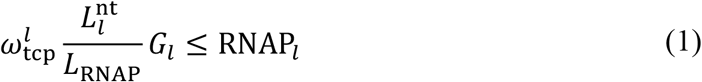

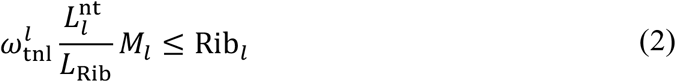

Equations 1 and 2 impose a basal level for the RNA polymerases (RNAP_*l*_) and ribosomes (Rib_*l*_) allocated to the template *l*. This basal level is defined based on the copy number of the gene *l* (*G*_*l*_) or the mRNA transcript *l* (*M*_*l*_), the footprint of RNA polymerase (*L*_RNAP_) or ribosome (*L*_Rib_) in nucleotides, the length of the template in nucleotides 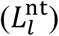, and the affinity of the RNA polymerase or ribosome for the template *l* reflected in 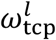and 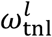, respectively. The constraints in Equations 1 and 2 can be defined for both native and heterologous genes. However, we applied Equations 1 and 2 only to the heterologous genes, as these genes are present in the host in high copy numbers due to the high copy number of plasmids. We assumed the basal level of RNA polymerases, and hence ribosomes, allocated to the native genes is negligible, as these genes are usually present in a single copy.

#### Allocation

In ETFL models, we divide the native proteins into two groups: (i) the ME-enzymes and (ii) the other proteins. The latter are not explicitly modeled in ETFL and are represented by a modeling protein called dummy protein. Then, we add a constraint of the following form to determine the fraction of the cellular protein content that can be allocated to the dummy protein (16):

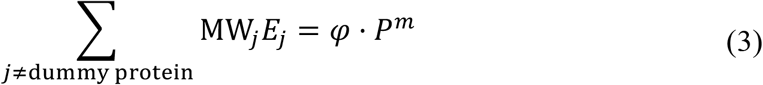

where MW_*j*_ and *E*_*j*_ are the molecular weight and molar concentration of *j*th protein, respectively. *P*^*m*^ is the fraction of the cell weight that is protein, and *φ* is the fraction of total protein allocated to the ME-enzymes. We used proteomics data to calculate this fraction as 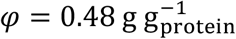. Since the total protein content *P*^*m*^ is fixed, Equation 3 also defines the share of dummy protein to be (1 − *φ*) · *P*^*m*^.

A part of the protein content is allocated to the heterologous proteins in a recombinant cell. However, since whole-cell proteomics data is not readily available for recombinant cells, it is difficult to determine the influence of recombinant proteins on *φ*. In the absence of proteomics data, we modified Equation 3 as follows:

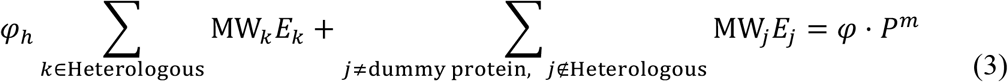

*φ*_*h*_ is a parameter representing the fraction of the heterologous proteins that take their share from the ME-enzymes (Figure 5).

**Figure 5:**
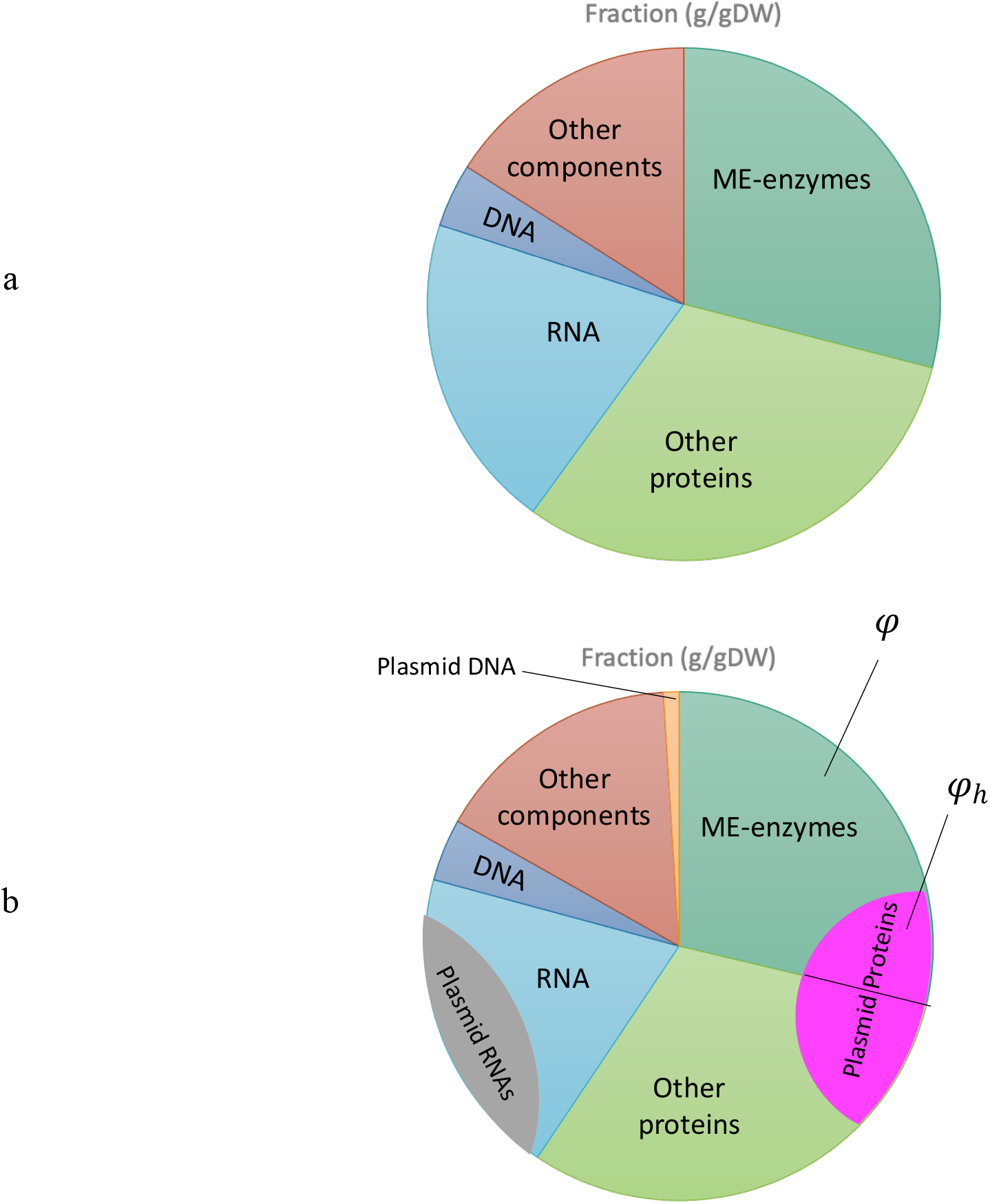
Schematic representation of the cellular composition. **a** wild-type cell and **b** recombinant cell. We assumed that apart from the plasmid DNA share, which increases due to the plasmid integration, the composition of the recombinant cell was the same as the wild-type cell. *φ* is a parameter representing the share of the total protein allocated to metabolism and expression. In the recombinant cell, the fractions of the cellular weight allocated to RNA and protein also include the heterologous RNAs and proteins, respectively. *φ*_*h*_ represents the fraction of the heterologous proteins taking their share from the metabolism- and expression-related enzymes.

### Estimation of 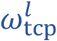and 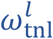

The parameters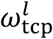and 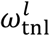represent the RNA polymerase and ribosome affinity for the gene and mRNA template *l*, respectively. Table S2 summarizes the fraction of RNA polymerases 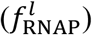and ribosomes 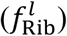allocated to plasmid-related expression. These fractions were calculated based on the available kinetic information. We varied 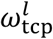and 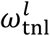 and solved the rETFL problem to calculate 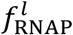and 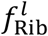subject to different plasmid copy numbers. Figure S1 shows that 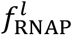 only depends on 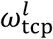, while Figure S2 shows that 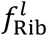 is impacted by variations in both 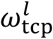 and 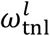. For each plasmid copy number, we selected 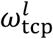and 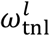 such that the following expression is minimized:

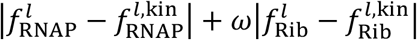

where 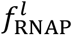and 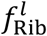are calculated by the rETFL problem, and 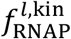and 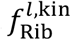are calculated using the kinetic parameters (Table S2). To check if the variation in *φ*_*h*_ impacts 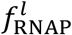and 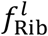, we calculated 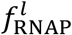 and 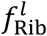subject to different *φ*_*h*_s. Figures S3 and S4 demonstrate that 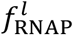 and 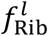 are independent of *φ*_*h*_.

### Estimation of *φ*_*h*_ and ATP maintenance

We used the experimental data for growth to estimate *φ*_*h*_ and ATPM. We varied *φ*_*h*_ and ATPM and maximized growth. We plotted the maximum growth rate for different values of *φ*_*h*_ and ATPM (Figure S5). At different copy numbers, changing ATPM had a uniform impact since ATPM was independent of the amount of heterologous protein production. However, the impact of *φ*_*h*_ was accentuated by increasing the copy number as more heterologous proteins were produced. That is, the growth reduction at low copy numbers depended on ATPM, and the slope of the reduction with increasing the copy number depended on *φ*_*h*_. We then chose ATPM and *φ*_*h*_ for which we obtained the best fit to the experimental data.

## Code Availability

The ecETFL model and the code used to create the models and perform the analyses is available at https://github.com/EPFL-LCSB/ecetfl.

## Acknowledgment

This work was funded by the European 528 Union’s Horizon 2020 research and innovation programme under grant agreement No 529 814408 and the Swiss National Science Foundation under grant agreement 530 200021_188623.

## Author Contribution

OO and VH conceptualized the study. OO adapted the code and ran the simulations. OO and VH discussed and visualized the results. OO and VH wrote the manuscript.

